# CRISPR/Cas12a toolbox for genomic manipulation in *Methanosarcina acetivorans*

**DOI:** 10.1101/2023.01.31.526428

**Authors:** Ping Zhu, Jichen Bao, Silvan Scheller

**Author notes:** **Corresponding authors:** Silvan Scheller, Jichen Bao, and.

## Abstract

Methanogenic archaea play an important role in the global carbon cycle and are regarded as promising host organisms for the biotechnological generation of fuels and chemicals from one-carbon substrates. *Methanosarcina acetivorans* is extensively studied as a model methanogen due to the availability of genetic tools and its versatile substrate range. Although genome editing in *M. acetivorans* via CRISPR/Cas9 has already been demonstrated, we now describe a user-friendly CRISPR/Cas12a toolbox that recognizes a T-rich (5′-TTTV) PAM sequence. This new system can manage deletions of 3500 bp (i.e., knockout of the entire *frhADGB* operon) and heterologous gene insertions with 80% efficiency observed in ten Pur^R^ transformants. Our CRISPR/Cas12a system also enables multiplex genome editing at high efficiency, which helps speed up genetic engineering. Deletions of 100 bp generated on two separate sites of the genome yielded 8/8 correctly edited transformants. Simultaneous gene deletion (100 bp) and replacement (100-bp region replaced by the 2400-bp *uidA* expression cassette) at a separate site was achieved, with 3/6 of transformants being edited correctly. In combination with the Cas9-based system, our CRISPR/Cas12a toolbox enables targeted genome editing at two sites (guanine-rich and thymine-rich, respectively) and, in so doing, hastens the overall genetic engineering of the *Methanosarcinales* species.

## Introduction

Methanogenic archaea strongly impact the global carbon cycle due to a contributive role in methane production (Kurth et al., 2020). Among the well-studied are the *Methanosarcina* species, as many details about their genomic information (Galagan et al., 2002; Maeder et al., 2006; Deppenmeier et al., 2002) and versatile methanogenesis (Costa and Leigh, 2014) have been earlier revealed. Owing to their efficient central metabolism (Carr and Buan, 2022), these methanogens allow for a wide range of metabolic applications (i.e., expanding substrates and energy sources, producing precursors for high-value products (McAnulty et al., 2017), and improving methane yields (Catlett et al., 2015)) through the rewiring of the methanogenesis pathway.

The CRISPR (clustered regularly interspaced short palindromic repeats) system is a powerful genetic editing tool and has already been widely developed for use in many eukaryotic and prokaryotic model organisms (Gao et al., 2002; Wang et al., 2021; Zhao et al., 2016). On the other hand, the application of CRISPR technology in archaeal species is still lagging far behind these other counterparts. In methanogens, the CRISPR system only came into use when the CRISPR/Cas9 system was first developed in *Methanosarcina acetivorans* (Nayak and Metcalf, 2017). Soon after, the CRISPRi-dCas9 tool was constructed for gene regulation in this same species (Dhamad and Lessner, 2020). More recently, CRISPR technology in another representative methanogenic species was reported, as the Cas12a- and Cas9-mediated CRISPR systems were developed in *Methanococcus maripaludis* (Bao et al., 2022; Li et al., 2022).

Cas12a (also known as Cpf1) is a class 2 type V endonuclease (Paul and Montoya, 2020) that, rather than recognizing a guanine (G)-rich (NGG-3′) protospacer adjacent motif (PAM) as with Cas9, will recognize a thymine (T)-rich (5′-TTTV, V□=□A, G, and C) PAM, thereby expanding the number of potential targeting sites along the genome. Moreover, the double-stranded breaks (DSB) caused by Cas12a will generate sticky ends, which help contribute to DNA repair and genome stability during genetic manipulation (Vanegas et al., 2019). As only the guide RNA (spacer) and direct repeat (DR) sequences are key elements of the Cas12a-based system, edited construction becomes simplified, particularly when multiplex genome editing is employed (Bandyopadhyay et al., 2020).

With an aim to further expand the CRISPR editing toolbox for genetic manipulation in *M. acetivorans*, we constructed the CRISPR/Cas12a system that expresses the Cas12a endonuclease from the *Lachnospiraceae bacterium* (Lb). To gauge the performance of this editing toolbox, single gRNA-mediated gene knockout and heterologous gene integration approaches were both evaluated. Finally, a crRNA (CRISPR RNA) array was constructed to establish the efficiency of Cas12a/crRNA-mediated multiplex genome editing, for which gene deletion and heterologous gene integration were obtained simultaneously at the two sites in the genome.

## Results

### Construction of Cas12a-gRNA expression system

To generate a Cas12a-gRNA expression system that is cloneable in most *Escherichia coli* strains, the *E. coli/Methanosarcina* shuttle vector pM000 (Supplementary Figure 1B) was constructed from the pWM321 plasmid (Metcalf et al., 1997) by first replacing the origin of replication (*ori*) from plasmid R6K with the one from plasmid ColE1. To establish markerless editability in the expression system, the hypoxanthine phosphoribosyltransferase (*hpt*) gene that confers sensitivity to the purine analog 8-aza-2,6-diaminopurine (8-ADP) for counter selection (Ehlers et al., 2011) was inserted downstream of the puromycin acetyltransferase (*pac*) gene, producing plasmid pM001 (Supplementary Figure 1C). The Cas12a cassette component, which consists of the Cas12a from *Lachnospiraceae bacterium* (Lb) amplified from pY016 (Zetsche et al., 2015) and fused to promoter PmcrB (tetO1), was inserted into the multiple cloning site (MCS) of pM001, producing plasmid pMCp4. The Cas12a-gRNA expression system (Figure 1A) was finalized by the construction and assembly of the gRNA cassette into pMCp4.

**Figure 1.**
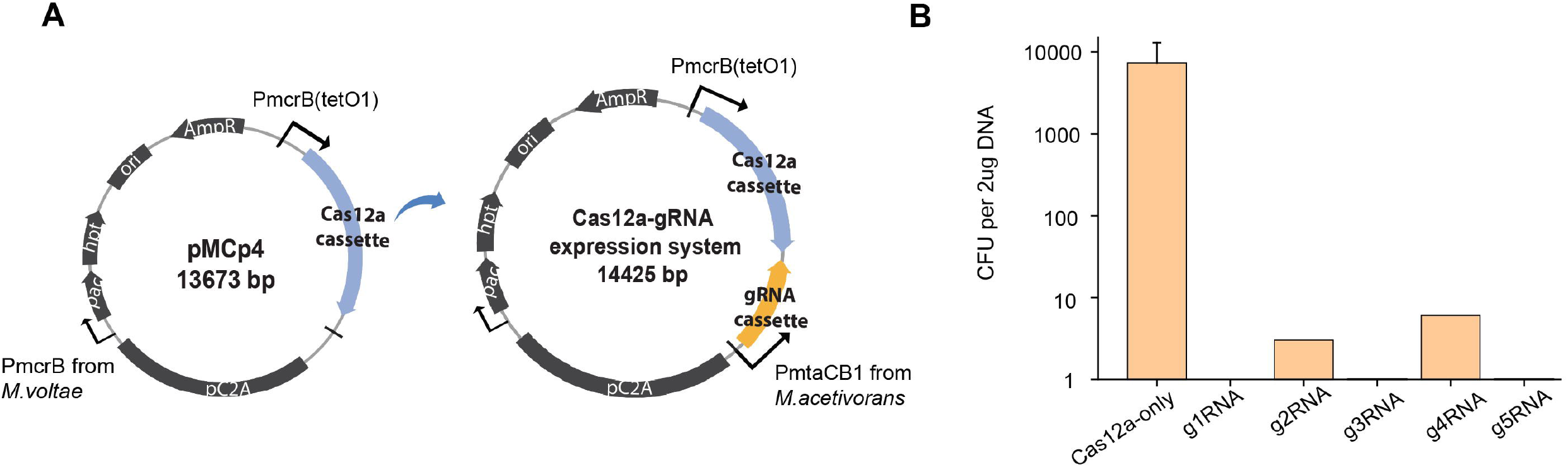
Construction of Cas12a-gRNA expression system for *M. acetivorans*. **(A)** Schematic diagram of pMCp4 and Cas12a-gRNA expression system. Plasmid pMCp4 expresses the Cas12a protein in *M. acetivorans*. Cas12a-gRNA expression system expresses the Cas12a-gRNA complex in *M. acetivorans*. Cas12a and gRNA cassettes are equipped with tetracycline-regulated promoter PmcrB (tetO1) and the promoter PmtaCB1 from *M. acetivorans*, respectively. **(B)** Targeting efficiency of Cas12a-gRNA expression system. Cas12a-only, plasmid pMCp4 that expresses Cas12a protein in *M. acetivorans*. g1RNA, g2RNA, g3RNA, g4RNA, and g5RNA, five gRNAs designed for targeting various locations along the *ssuC* gene. Error bar represents the standard deviation of triplicate measurements. Standard deviations were not determined for gRNA-expressing transformation data, as all cells were plated out to analyze the lethal efficiency of the Cas12a-gRNA complex.

The possible cytotoxicity of the Cas12a protein was determined by transforming the shuttle vector pM001 and the Cas12a-expressing plasmid pMCp4 into *M. acetivorans* and comparing the efficiency rates. When 2 μg amounts of plasmid DNA were used to transform *M. acetivorans*, pM001 yielded 40,000 ± 20,000 CFU of Pur^R^ transformants, whereas pMCp4 yielded 34,000 ± 19,000 CFU of Pur^R^ transformants (Supplementary Figure 2A). As no significant difference (two-tailed *t-test, P* = 0.75) was observed in the transformation efficiencies, this indicates that the expression of Cas12a protein is non-toxic and not harmful to *M. acetivorans* cells. Growth curve analysis of *M. acetivorans* suggests that the presence of pM001 and pMCp4 place no growth pressure on cells (Supplementary Figure 2B), thus allowing the use of both plasmids in genome editing applications.

To evaluate the targeting efficiency of Cas12a-gRNA expression system, five different gRNAs were used to target various locations within the non-essential *ssuC* gene (which encodes the permease subunit of the sulfonate ABC transporter) in the genome. Less than 10 CFUs (per 2 μg DNA) were observed from the cells expressing the Cas12a-gRNA complex (Figure 1B), thus indicating the high targeting efficiency of all five gRNAs and the strength of the constructed system.

### CRISPR/Cas12a-mediated gene knockout in *M. acetivorans*

To investigate the efficiency of gene deletion, the homologous repair (HR) arms for genome editing were inserted into the Cas12a-gRNA expression system to produce the Cas12a-gRNA genome editing system (Figure 2A). The *ssuC* gene was targeted for knockout. Here, the g1RNA was used to target downstream of the *ssuC* gene. For gRNA cassette construction, PCR amplification was used to synthesize the gRNA and DR sequences that contain promoter (PmtaCB1) and terminator (TmtaCB1) regions (Supplementary Figure 3A). Deletion lengths varied from 100 to 2000 bp (Figure 2B) and a 1000-bp length of flanking HR sequence was chosen as optimal based on previous work (Nayak and Metcalf, 2017).

**Figure 2.**
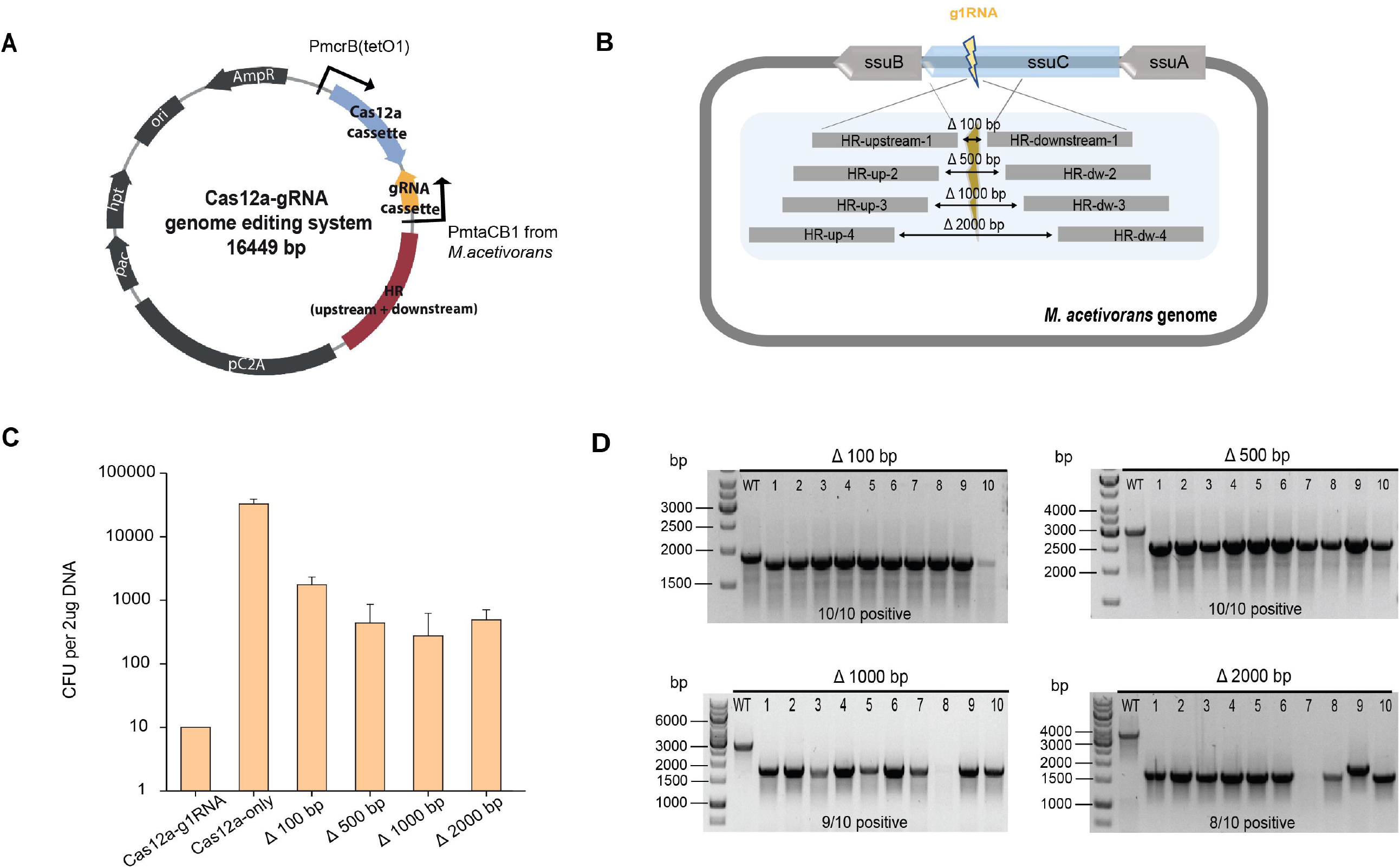
CRISPR/Cas12a-mediated gene knockout in *M. acetivorans*. **(A)** Schematic diagram of Cas12a-gRNA genome editing system. Cas12a and gRNA cassettes are equipped with promoters PmcrB (tetO1) and PmtaCB1, respectively. Genome editing was achieved by the introduced upstream and downstream homologous repair (HR) arms. **(B)** Scheme for generating gene deletions in *ssuC*. g1RNA was designed to target downstream of *ssuC* for generating a double-stranded break (DSB). Various sizes of gene knockouts were generated by introducing a 1000-bp length of flanking HR sequence near the leakage site. Δ100 bp, Δ500 bp, Δ1000 bp, and Δ2000 bp, plasmids generating 100 bp, 500 bp, 1000 bp, and 2000 bp gene knockouts while repairing the leakage. **(C)** Transformation efficiency of deletion-generating plasmids. Cas12a-g1RNA, plasmid expressing Cas12a-g1RNA complex that targets *ssuC* to produce a DSB on the genome. Cas12a-only, plasmid pMCp4 expresses the Cas12a protein. Error bars represent the standard deviation of triplicate measurements. Standard deviations were not determined for Cas12a-g1RNA transformation data, as all cells were plated out to analyze the lethal efficiency of the Cas12a-g1RNA complex. **(D)** The editing efficiency of deletion-generating plasmids. Ten Pur^R^ transformants were randomly selected for colony PCR. WT, wide type *M. acetivorans* strain. Thermo Scientific™ GeneRuler 1kb DNA ladder was used for sizing DNA fragments.

When pMCp4 (only Cas12a) was transformed into *M. acetivorans*, this yielded a high number of Pur^R^ transformants (33,000 ± 6,000 CFU per 2 μg DNA). On the other hand, only ten Pur^R^ transformants were observed in cells transformed with plasmid pMCp2-g1RNA (see Table 2), where the Cas12a-g1RNA complex was being expressed (Figure 2C). The 100-bp deletion from the genome had the highest repairing efficiency (1,800 ± 600 CFU) (Figure 2C). This is consistent with the previous results of a Cas9-mediated gene knockout study, in which shorter deletions (i.e., Δ 100 bp and Δ 500 bp) were considered more stable and reliable (Nayak and Metcalf, 2017). For the plasmids that led to longer deletions (i.e., Δ1000 bp and Δ2000 bp), similar transformant yields were obtained (Figure 2C). The knockout (or editing) efficiency of the deletion-generating plasmids was verified for ten transformants by colony PCR (Figure 2D) using the primers listed in Table 3. A higher editing efficiency was observed with the shorter deletions, in which a 100% of the 100 bp-deleted transformants were positive. Interestingly though, 80% of the 2000 bp-deleted transformants were also edited positively. In addition, three samples from each transformation set had their genomes sequenced, with all genomes being edited correctly (data not shown).

**Table 1.** Strains used in this study.

| Strain | Properties | Source |
| --- | --- | --- |
| *Methanosarcina acetivorans* |  |  |
| WWM73 | Δhpt::PmcrB-tetR-ϕC31-int-attP | (Guss et al., 2008) |
| M73-100 | Δhpt::PmcrB-tetR-ϕC31-int-attP, ΔssuC (72928-73101)::GCGGCCGC | This study |
| M73-500 | Δhpt::PmcrB-tetR-ϕC31-int-attP, ΔssuC (72782-73301)::GCGGCCGC | This study |
| M73-1000 | Δhpt::PmcrB-tetR-ϕC31-int-attP, ΔssuB ΔssuC (72532-73551)::GCGGCCGC | This study |
| M73-2000 | Δhpt::PmcrB-tetR-ϕC31-int-attP, ΔssuB ΔssuC ΔssuA (72032-74051)::GCGGCCGC | This study |
| M73-3500 | Δhpt::PmcrB-tetR-ϕC31-int-attP, ΔfrhADGB (1165689-1169233)::GCGGCCGC | This study |
| M73-100-100 | Δhpt::PmcrB-tetR-ϕC31-int-attP, ΔssuC (72928-73101)::GCGGCCGC, ΔfrhA (1166033-1166145)::GCGGCCGC | This study |
| M73-uid | Δhpt::PmcrB-tetR-ϕC31-int-attP, ΔssuC (72928-73101)::Pmcr-uidA-Tmcr | This study |
| M73-uid-100 | Δhpt::PmcrB-tetR-ϕC31-int-attP, ΔssuC (72928-73101)::Pmcr-uidA-Tmcr, ΔfrhA (1166033-1166145)::GCGGCCGC | This study |
| *Escherichia coli* |  |  |
| DH10B | F<sup>-</sup> mcrA $\Delta$(mrr-hsdRMS-mcrBC) ϕ80lacZΔM15 ΔlacX74 recA1 endA1 araD139 $\Delta$(ara-leu)7697 galU galK λ- rpsL (StrR) nupG | New England Biolabs |
| XL10-Gold | TetrD (mcrA)183 D (mcrCB-hsdSMR-mrr) 173 endA1 supE44 thi-1 recA1 gyrA96 relA1 lac Hte [F' proAB lacIqZDM15 Tn10 (Tetr) Amy Camr] | Agilent Technologies |

**Table 2.** Plasmids used in this study.

| Plasmid | Description | Source |
| --- | --- | --- |
| pY016 | gene sequence source of LbCas12a | (Zetsche et al., 2015) |
| pWM321 | *Escherichia coli*/*Methanosarcina* shuttle vector | (Metcalf et al., 1997) |
| pM000 | pWM321-derived vector, ColE1 origin of replication (high-copy-number from *E.coli*) | This study |
| pM001 | pM000-derived vector with hypoxanthine phosphoribosyltransferase (*hpt*) gene | This study |
| pMCp4 | pM001-derived plasmid with Cas12a cassette | This study |
| pMCp2-g1RNA | pMCp4-derived plasmid with gRNA cassette, targeting *ssuC* locus | This study |
| pMCp3-g1-100 | pMCp2-g1RNA-derived plasmid with flanked homologous repair arms generating 100 bp of deletion on genome | This study |
| pMCp3-g1-500 | pMCp2-g1RNA-derived plasmid with flanked homologous repair arms generating 500 bp of deletion on genome | This study |
| pMCp3-g1-1000 | pMCp2-g1RNA-derived plasmid with flanked homologous repair arms generating 1000 bp of deletion on genome | This study |
| pMCp3-g1-2000 | pMCp2-g1RNA-derived plasmid with flanked homologous repair arms generating 2000 bp of deletion on genome | This study |
| pMCp2-g9RNA | pMCp4-derived plasmid with gRNA cassette, targeting *frhA* locus |  |
| pMCp3-g9-3500 | pMCp2-g9RNA-derived plasmid with flanked homologous repair arms generating 3500 bp of deletion on genome | This study |
| pMCp3-g1-100-uid | pMCp2-g1RNA-derived plasmid with flanked homologous repair arms with *uidA* expression cassette inserted in genome | This study |
| pMCp2-g1g9RNA | pMCp4-derived plasmid with gRNA expression cassette, targeting *ssuC* and *frhA* locus simultaneously | This study |
| pMCp3-g1g9-100 | pMCp2-g1g9RNA-derived plasmid with two sets of homologous repair arms generating 100 bp deletions on *ssuC* and *frhA* locus simultaneously | This study |
| pMCp3-g1-uid-g9-100 | pMCp2-g1g9RNA-derived plasmid with two sets of homologous repair arms with *uidA* expression cassette inserted in *ssuC* locus and 100 bp deletion in *frhA* locus simultaneously | This study |

### CRISPR/Cas12a-mediated gene insertion in *M. acetivorans*

In order to assess gene insertion by the Cas12a-gRNA genome editing system, a heterologous gene cassette was placed within the HR sequences and utilized for integrating the β-glucuronidase (*uidA*) gene, given its previous successful use in *Methanosarcina* (Guss et al., 2008). For *uidA* cassette construction, the *uidA* gene was PCR amplified from *E*.*coli* BL21 genomic DNA and fused to the mcr promoter (Pmcr) and terminator (Ter) sequences from *M. barkeri*. The insertion efficiency of the editing plasmid pMCp3-g1-100-uid (Δ100 bp:: *uidA*) (Figure 3A) is based on the number of Pur^R^ transformants (1,700 ± 900 CFU per 2 μg DNA) obtained. The insertion efficiency was also shown for ten transformants by colony PCR, with 80% proving to be edited positively (Figure 3B).

**Figure 3.**
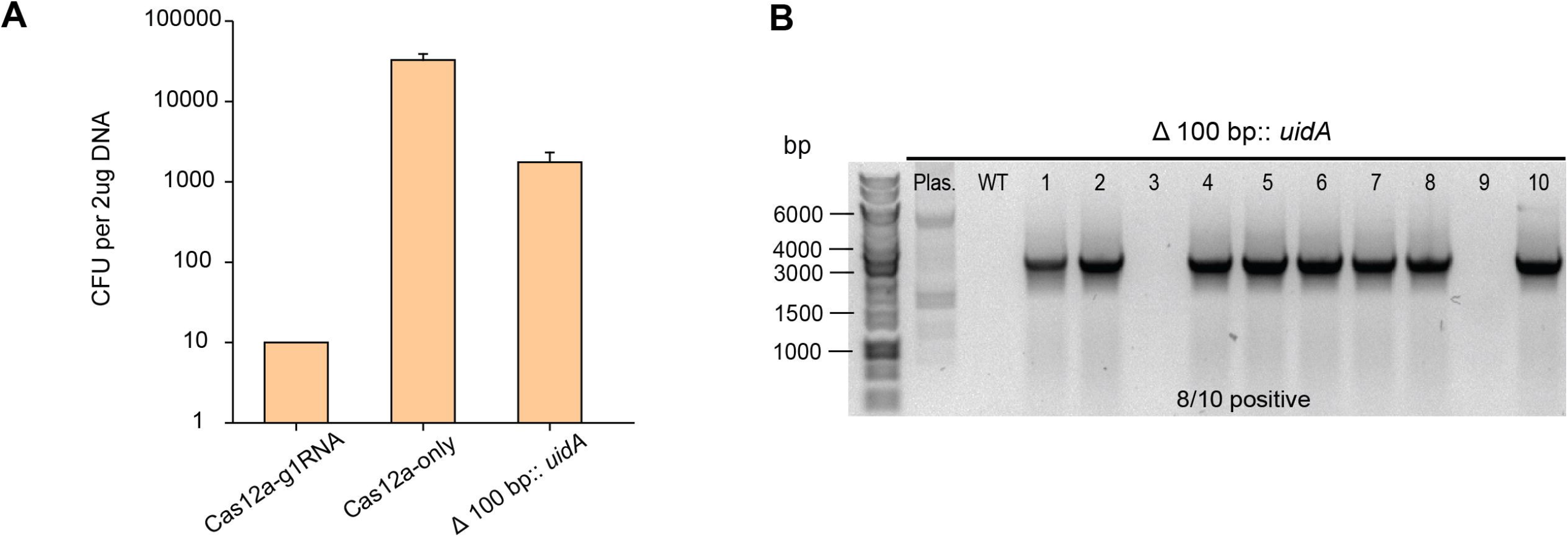
CRISPR/Cas12a-mediated gene insertion in *M. acetivorans*. **(A)** Transformation efficiency of Cas12a-mediated gene insertion. Cas12a-gRNA and Cas12a-only originate from the same transformation as gene knockouts. Δ100 bp:: *uidA* from two repeated transformations, the Cas12a-mediated editing system replaced a 100 bp region with the *uidA* cassette. Error bars represent the standard deviation of triplicate measurements. Standard deviations were not determined for Cas12a-g1RNA transformation data, as all cells were plated out to analyze the lethal efficiency of the Cas12a-g1RNA complex. **(B)** Ten Pur^R^ transformants were randomly selected for colony PCR to verify the insertion of the *uidA* gene in the genome. Plas. (plasmid pMCp3-g1-100-uid) and WT (wild type *M. acetivorans* genome) served as negative controls in the colony PCR. Thermo Scientific™ GeneRuler 1kb DNA ladder was used for sizing DNA fragments.

### Cas12a/crRNA-mediated multiplex genome editing in *M. acetivorans*

Cas12a/crRNA-mediated multiplex genome editing was carried out using the pMCp2-g1g9RNA plasmid. Here, two gRNA sequences (g1RNA and g9RNA), both equipped with promoter PmtaCB1 and terminator TmtaCB1, were used to simultaneously edit the *ssuC* and *frhA* (which encodes the coenzyme F420 hydrogenase alpha subunit) genes (Figure 4A and Supplementary Figure 4). For assessing the effect of double-site deletions, two sets of HR sequences that generate 100-bp deletions in each of the genes (see above) were designed and assembled into pMCp2-g1g9RNA, producing the plasmid pMCp3-g1g9-100. Among the obtained transformants (107 ± 23 CFU per 2 μg DNA), eight were randomly selected for PCR amplification to corroborate the editing effect. Two pairs of primers (veri1/veri2 and veri11/Cp54) were used to separately target upstream and downstream of the leakage sites in the *ssuC* and *frhA* genes. All eight transformants had proved to be positive (Figure 4B). For assessing the versatility of multiplex genome editing, another plasmid (pMCp3-g1-uid-g9-100) was constructed that inserts the *uidA* cassette into *ssuC* as well as makes a 100-bp deletion in *frhA* during a single transformation (Figure 4C). Six Pur^R^ transformants were randomly selected for separate colony PCR verification of the *ssuC* and *frhA* sites using two pairs of primers (veri2/veri8 and veri9/veri10). Here, distinctive editing effects were observed, as only 50% of transformants had contained the *uidA* cassette within the *ssuC* gene, while more the 80% of transformants showed a 100-bp deletion within the *frhA* gene (Figure 4D).

**Figure 4.**
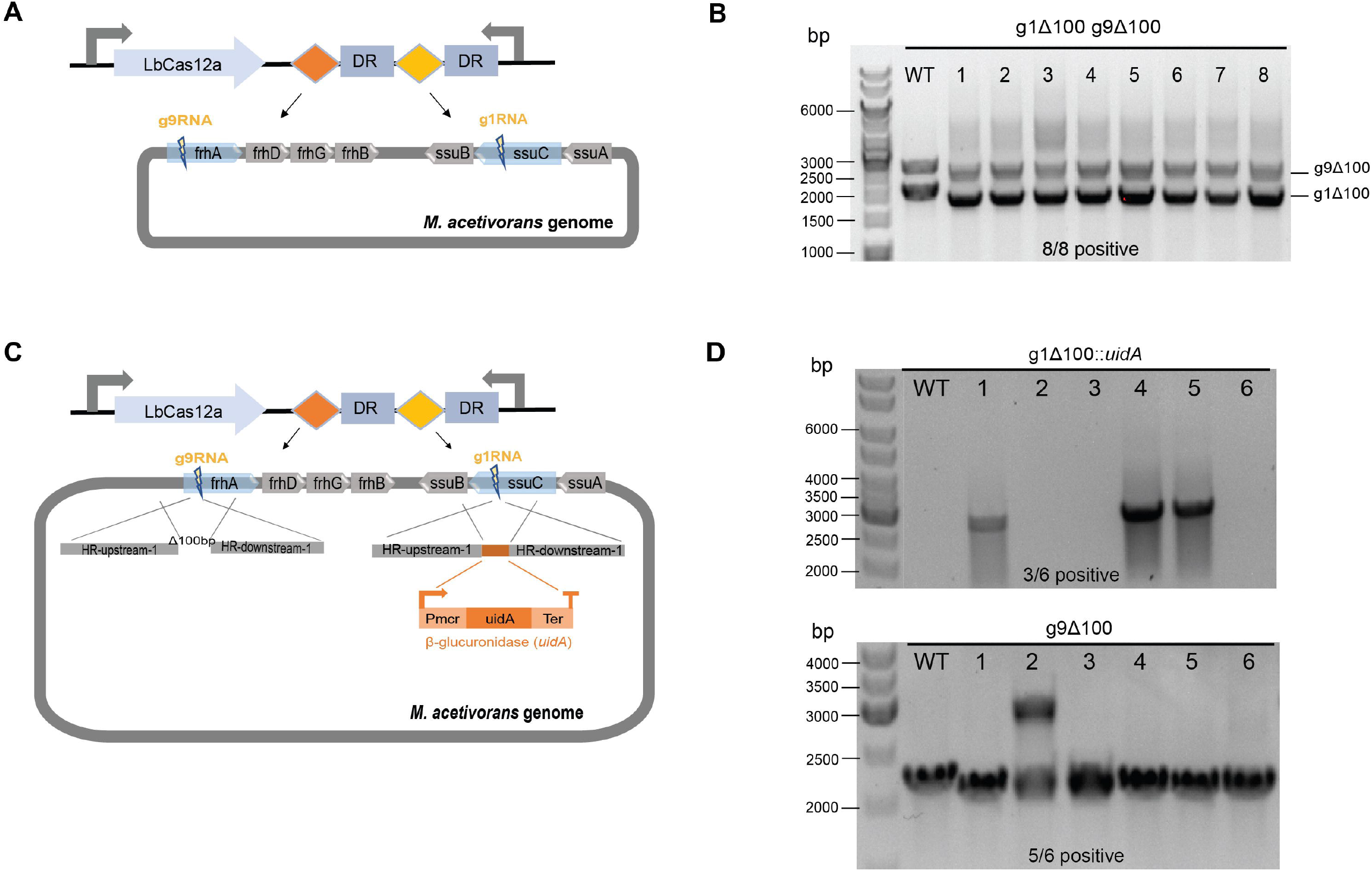
CRISPR/Cas12a-mediated multiplex genome editing in *M. acetivorans*. **(A)** crRNA array designed for targeting two sites on the genome. g1RNA and g9RNA target *ssuC* and *frhA*, generating two leakages on the genome simultaneously. DR, direct repeat sequences in the gRNA cassette. **(B)** Efficiency of multiplex gene knockout. 100 bp of gene sequences in *ssuC* and *frhA* were deleted simultaneously. Eight transformants were randomly selected for colony PCR to verify the gene knockout efficiency. WT, wide type *M. acetivorans* strain served as the negative control. g1Δ100 bp, gene deletion in *ssuC*. g9Δ100 bp, gene deletion in *frhA*. Thermo Scientific™ GeneRuler 1kb DNA ladder was used for sizing DNA fragments. **(C)** Scheme for Cas12a-mediated simultaneous gene insertion and deletion. g1RNA-interfered targeting replaces 100 bp region with the *uidA* cassette in *ssuC* and g9RNA-interfered targeting generates 100 bp deletion in *frhA*. **(D)** Efficiency of the simultaneous gene insertion and deletion. g1Δ100 bp::*uidA*, gene replacement in *ssuC*. g9Δ100 bp, gene deletion in *frhA*. WT, wide type *M. acetivorans* strain served as the negative control. Thermo Scientific™ GeneRuler DNA Ladder Mix was used for sizing DNA fragments.

## Discussion

Spurred on by the development of the innovative CRISPR-mediated editing system, building and optimizing a new generation of CRISPR tools for different organisms is emerging as a leading strategy in many research areas. In our study, we developed the CRISPR/Cas12a system to expand the genetic engineering toolbox for *M. acetivorans*. Unlike the CRISPR/Cas9 system, our Cas12a-based CRISPR tool recognizes T-rich regions along the genome as well as edits using a more simplified mechanism that does not rely on tracrRNA (trans-activating CRISPR RNA) for crRNA maturation (Nakade et al., 2017). Here, Cas12a-mediated genome editing proved to be very effective for performing gene deletions and heterologous gene insertions. Moreover, multiplex genome editing of *M. acetivorans* using our CRISPR/Cas12a system had clearly demonstrated the design simplicity of the crRNA array and the robust potential for multifunctional editing using only a single plasmid construct.

Owing to the possible cytotoxicity of the Cas endonuclease, cellular expression of CRISPR/Cas systems can occasionally be a detriment and give low transformation yields (Wendt et al., 2016). Since no harmful effects were observed with LbCas12a protein expression in *M. acetivorans*, this indicates that our CRISPR/Cas12a tool is a suitable approach for further genome editing. Achieving a higher targeting efficiency usually requires the simultaneous screening of multiple gRNA sequences, and then often entails using laborious and multi-step methods to construct several plasmids. As a novel simplification, we constructed plasmid pMCp2-gX (Supplementary Figure 6), which allows for quick preparation (10 mins) of the gRNA sequences and their seamless assembly into linearized vectors for the efficient generation of multiple Cas12a-gRNA-expressing plasmids.

Typically, many factors can affect the editing performance of CRISPR/Cas systems, such asthe genome regions to be edited, gRNA targeting efficiency and precision, and HR sequence length. Performing these genetic manipulations can often be problematic in *M. acetivorans*, particularly if little is known beforehand about a specific targeted genome region. For the present study, the *ssuC* and *frhA* genes were selected as the target sites in the genome since their protein products are non-essential in *M. acetivorans* (Nayak and Metcalf, 2017; Guss et al., 2009). Also, a 23-nt gRNA was empirically chosen, as this sequence length displayed an improved targeting specificity that was higher than a shorter 20-nt gRNA (data not shown). Conversely, shorter gRNA sequence lengths in the Cas9-mediated system proved to be more effective (Fu et al., 2014). In order to promote homologous repair of the double-stranded breaks, a flanking homology arm of 1000 bp was used with all genome edits, as this particular length exhibited a high repairing efficiency in *M. acetivorans* and *M. maripaludis* (Bao et al., 2022).

By comparison, the generation of long fragment deletions are reported to be more efficient with the CRISPR/Cas12a system than the Cas9-based one (Wang et al., 2022). Similarly, our study also showed that Cas12a-mediated long fragment deletion of 2000 bp yielded a higher number of transformants than obtained with CRISPR/Cas9 editing (Nayak and Metcalf, 2017). To establish the possible length limits of gene deletion with our CRISPR/Cas12a system, the plasmid pMCp3-g9-3500 was constructed for removing the entire *frhADGB* operon, which resulted in a 3500 bp gene knockout from the genome. Here, a higher number of Pur^R^ transformants (2,800 ± 1,400 CFU) were obtained, and among the ten randomly selected for PCR verification 100% proved to be positive (Supplementary Figure 5). This suggests that our CRISPR/Cas12a editing tool is highly efficient for making long fragment deletions.

In this present study, multiplex genome editing with the Cas12a/crRNA array was assessed to shorten the genetic engineering process in *M. acetivorans*. Our data revealed that this system has great potential for simultaneously generating gene deletions and insertions, albeit a lower number of transformants were obtained than with single gRNA editing. Also, it now appears possible to design the crRNA array comprised of additional gRNAs for multiple site-targeting in *M. acetivorans*, given the previous study showed that four genomic loci were successfully edited in mammalian cells using a single crRNA array, and with no editing dominance among the various gRNAs (Zetsche et al., 2017).

Irrespective of gene deletion or insertion, similarly high performances were achieved with single-site genome editing as at least 8/10 of the randomly selected Pur^R^ transformants had been edited positively. On the other hand, these results appeared to vary after plasmid curing for five randomly selected transformants since 100% of 8ADP^R^ transformants exhibited the 100-bp gene deletion, whereas only 40% of 8ADP^R^ transformants had a correctly inserted *uidA* cassette (Supplementary Figure 7). This implies that the editing performance is distinctive if plasmid curing is involved.

Likewise, with the multiplex genome edits, the *uidA* gene insertion within the *frhA* site occurred less efficiently than the 100-bp deletion of the *ssuC* gene. Nonetheless, it is well worth noting that since the β-glucuronidase (*uidA*) reporter strategy is routinely used in *M. acetivorans* (Guss et al., 2009), this protein should pose no harm to growing transformants. It thus appears that the *M. acetivorans* cells edited with only gene deletions will likely be more stable than those with gene insertions involved.

To conclude, our CRISPR/Cas12a editing tool can efficiently implement the genomic manipulation of *M. acetivorans*, particularly long fragment deletions. In addition, the unique mechanism of the Cas12a endonuclease also streamlines the crRNA array design for multiplex genome editing, potentially enabling a broad range of new genetic engineering applications.

## Material and methods

### Strains and media

High-salt (HS) broth medium containing 125 mM methanol was used for cultivating all *M. acetivorans* strains at 37°C. *M. acetivorans* derivatives in this study are listed in Table 1. *M. acetivorans* WWM73 was used as the host strain in this study. HS solid medium containing 1.4% agar and 2 μg/mL puromycin (InvivoGen, Inc.) was used for screening *M. acetivorans* transformants. *E*.*coli* DH10B (Thermo Scientific) or *E*.*coli* XL10-Gold (Agilent Technologies) was used as the cloning host for constructing CRISPR plasmids. *E. coli* transformants were selected using lysogeny broth (LB) solid medium supplemented with 100 μg/mL ampicillin.

### Primers and plasmid construction

All plasmids used in this study are listed in Table 2. All primers are listed in Table 3. PrimeSTAR^®^ Max DNA Polymerase (Takara Bio) was used for amplifying gene fragments by PCR. SapphireAmp Fast PCR Master Mix (Takara Bio) was used for verifying *E*.*coli* and *M. acetivorans* transformants by colony PCR. HiFi DNA Assembly Master Mix (New England BioLabs) was used for Gibson assembly of DNA unless otherwise stated. crRNA sequences for multiplex genome targeting were synthesized and combined using the Gibson assembly method. Donor DNA with upstream and downstream homology arms was inserted immediately downstream of the gRNA cassette to construct the CRISPR/Cas12a genome editing system, otherwise known as plasmid pMCp3-X (see Table 2).

### DNA transformation methods

Polyethylene glycol (PEG)-mediated transformation of *M. acetivorans* was performed with 2 μg DNA according to the method described previously (Oelgeschläger and Rother, 2009). *M. acetivorans* transformant cells were grown on plates of HS solid medium and incubated at 37°C for 10-15 days in an anaerobic jar with a controlled headspace gas mix of 79.95% N_2_, 20% CO_2_, and 0.05% /H_2_S. Chemically competent *E*.*coli* cells were used for the transformation of assembled plasmids.

## Data Availability Statement

The original contributions presented in the study are included in the article/Supplementary Material, further inquiries can be directed to the corresponding authors.

## Author Contributions

P.Z., J.B. and S.S. conceived the study. P.Z. performed the experiments and wrote the manuscript. J.B. and S.S. supervised the experiments and revised the manuscript.

All authors declare no conflict of interest.

## Acknowledgments

We thank the Novo Nordisk Foundation (grant NNF19OC0054329 to S. S.; grant NNF20OC0065032 to J. B.) and the Academy of Finland (grant 326020 to S. S.) for funding this research. We thank the AScI project student Shreyash Borkar for helping with the plasmid transformation. We thank Prof. Michael Rother and Dr. Christian Schöne for testing the editing efficiency of the CRISPR/Cas12a system. We thank Dr. Ingemar von Ossowski for manuscript polishing and grammar checking.

## References

Bandyopadhyay, A., Kancharla, N., Javalkote, V. S., Dasgupta, S., and Brutnell, T. P. (2020). CRISPR-Cas12a (Cpf1): A Versatile Tool in the Plant Genome Editing Tool Box for Agricultural Advancement. Frontiers in Plant Science 11. doi: 10.3389/fpls.2020.584151.

Bao, J., De Dios Mateos, E., and Scheller, S. (2022). Efficient CRISPR/Cas12a-Based Genome-Editing Toolbox for Metabolic Engineering in Methanococcus maripaludis. ACS Synthetic Biology 11, 2496–2503. doi: 10.1021/acssynbio.2c00137.

Carr, S., and Buan, N. R. (2022). Insights into the biotechnology potential of Methanosarcina. Frontiers in Microbiology 13. Available at: https://www.frontiersin.org/articles/10.3389/fmicb.2022.1034674 [Accessed January 6, 2023].

Catlett, J. L., Ortiz, A. M., and Buan, N. R. (2015). Rerouting Cellular Electron Flux To Increase the Rate of Biological Methane Production. Appl Environ Microbiol 81, 6528–6537. doi: 10.1128/AEM.01162-15.

Costa, K. C., and Leigh, J. A. (2014). Metabolic versatility in methanogens. Current Opinion in Biotechnology 29, 70–75. doi: 10.1016/j.copbio.2014.02.012.

Deppenmeier, U., Johann, A., Hartsch, T., Merkl, R., Schmitz, R. A., Martinez-Arias, R., et al. (2002). The Genome of Methanosarcina mazei: Evidence for Lateral Gene Transfer Between Bacteria and Archaea. Available at: www.caister.com/bacteria-plant.

Dhamad, A. E., and Lessner, D. J. (2020). A CRISPRi-dCas9 System for Archaea and Its Use To Examine Gene Function during Nitrogen Fixation by Methanosarcina acetivorans. Applied and Environmental Microbiology 86. doi: 10.1128/AEM.01402-20.

Ehlers, C., Jäger, D., and Schmitz, R. A. (2011). Establishing a markerless genetic exchange system for Methanosarcina mazei strain Gö1 for constructing chromosomal mutants of small RNA genes. Archaea 2011, 1–7. doi: 10.1155/2011/439608.

Fu, Y., Sander, J. D., Reyon, D., Cascio, V. M., and Joung, J. K. (2014). Improving CRISPR-Cas nuclease specificity using truncated guide RNAs. Nature Biotechnology 32, 279–284. doi: 10.1038/nbt.2808.

Galagan, J. E., Nusbaum, C., Roy, A., Endrizzi, M. G., Macdonald, P., FitzHugh, W., et al. (2002). The Genome of M. acetivorans Reveals Extensive Metabolic and Physiological Diversity. Genome Res 12, 532–542. doi: 10.1101/gr.223902.

Gao, J., Xu, J., Zuo, Y., Ye, C., Jiang, L., Feng, L., et al. (2002). Synthetic Biology Toolkit for Marker-Less Integration of Multigene Pathways into Pichia pastoris via CRISPR/Cas9. doi: 10.1021/acssynbio.1c00307.

Guss, A. M., Kulkarni, G., and Metcalf, W. W. (2009). Differences in Hydrogenase Gene Expression between Methanosarcina acetivorans and Methanosarcina barkeri. Journal of Bacteriology 191, 2826–2833. doi: 10.1128/JB.00563-08.

Guss, A. M., Rother, M., Zhang, J. K., Kulkkarni, G., and Metcalf, W. W. (2008). New methods for tightly regulated gene expression and highly efficient chromosomal integration of cloned genes for Methanosarcina species. Archaea 2, 193–203. doi: 10.1155/2008/534081.

Kurth, J. M., Op den Camp, H. J. M., and Welte, C. U. (2020). Several ways one goal—methanogenesis from unconventional substrates. Applied Microbiology and Biotechnology 104, 6839–6854. doi: 10.1007/s00253-020-10724-7.

Li, J., Zhang, L., Xu, Q., Zhang, W., Li, Z., Chen, L., et al. (2022). CRISPR-Cas9 Toolkit for Genome Editing in an Autotrophic CO 2 -Fixing Methanogenic Archaeon. Microbiology Spectrum 10. doi: 10.1128/spectrum.01165-22.

Maeder, D. L., Anderson, I., Brettin, T. S., Bruce, D. C., Gilna, P., Han, C. S., et al. (2006). The Methanosarcina barkeri genome: Comparative analysis with Methanosarcina acetivorans and Methanosarcina mazei reveals extensive rearrangement within methanosarcinal genomes. Journal of Bacteriology 188, 7922–7931. doi: 10.1128/JB.00810-06.

McAnulty, M. J., Poosarla, V. G., Li, J., Soo, V. W. C., Zhu, F., and Wood, T. K. (2017). Metabolic engineering of Methanosarcina acetivorans for lactate production from methane. Biotechnology and Bioengineering 114, 852–861. doi: 10.1002/bit.26208.

Metcalf, W. W., Zhang, J. K., Apolinario, E., Sowers, K. R., and Wolfe, R. S. (1997). A genetic system for Archaea of the genus Methanosarcina□: Liposome-mediated transformation and construction of□shuttle□vectors. Proceedings of the National Academy of Sciences 94, 2626–2631. doi: 10.1073/pnas.94.6.2626.

Nakade, S., Yamamoto, T., and Sakuma, T. (2017). Cas9, Cpf1 and C2c1/2/3―What’s next? Bioengineered 8, 265–273. doi: 10.1080/21655979.2017.1282018.

Nayak, D. D., and Metcalf, W. W. (2017). Cas9-mediated genome editing in the methanogenic archaeon Methanosarcina acetivorans. Proceedings of the National Academy of Sciences of the United States of America 114, 2976–2981. doi: 10.1073/pnas.1618596114.

Oelgeschläger, E., and Rother, M. (2009). In vivo role of three fused corrinoid/methyl transfer proteins in Methanosarcina acetivorans. Molecular Microbiology 72, 1260–1272. doi: 10.1111/j.1365-2958.2009.06723.x.

Paul, B., and Montoya, G. (2020). CRISPR-Cas12a: Functional overview and applications. Biomedical Journal 43, 8–17. doi: 10.1016/j.bj.2019.10.005.

Vanegas, K. G., Jarczynska, Z. D., Strucko, T., and Mortensen, U. H. (2019). Cpf1 enables fast and efficient genome editing in Aspergilli. Fungal Biology and Biotechnology 6. doi: 10.1186/s40694-019-0069-6.

Wang, X., Dong, K., Kong, D., Zhou, Y., Yin, J., Cai, F., et al. (2021). A far-red light – inducible CRISPR-Cas12a platform for remote-controlled genome editing and gene activation. 2358, 1–12.

Wang, Z., Wang, Y., Wang, Y., Chen, W., and Ji, Q. (2022). CRISPR/Cpf1-Mediated Multiplex and Large-Fragment Gene Editing in Staphylococcus aureus. ACS Synthetic Biology. doi: 10.1021/acssynbio.2c00248.

Wendt, K. E., Ungerer, J., Cobb, R. E., Zhao, H., and Pakrasi, H. B. (2016). CRISPR/Cas9 mediated targeted mutagenesis of the fast growing cyanobacterium Synechococcus elongatus UTEX 2973. Microbial Cell Factories 15, 1–8. doi: 10.1186/s12934-016-0514-7.

Zetsche, B., Gootenberg, J. S., Abudayyeh, O. O., Slaymaker, I. M., Makarova, K. S., Essletzbichler, P., et al. (2015). Cpf1 Is a Single RNA-Guided Endonuclease of a Class 2 CRISPR-Cas System. Cell 163, 759–771. doi: 10.1016/j.cell.2015.09.038.

Zetsche, B., Heidenreich, M., Mohanraju, P., Fedorova, I., Kneppers, J., Degennaro, E. M., et al. (2017). Multiplex gene editing by CRISPR-Cpf1 using a single crRNA array. Nature Biotechnology 35, 31–34. doi: 10.1038/nbt.3737.

Zhao, D., Yuan, S., Xiong, B., Sun, H., Ye, L., Li, J., et al. (2016). Development of a fast and easy method for Escherichia coli genome editing with CRISPR/ Cas9. Microb Cell Fact 15, 205. doi: 10.1186/s12934-016-0605-5.

