## Supplementary figure 1-7 for "CRISPR/Cas12a toolbox for genomic manipulation in *Methanosarcina acetivorans*"

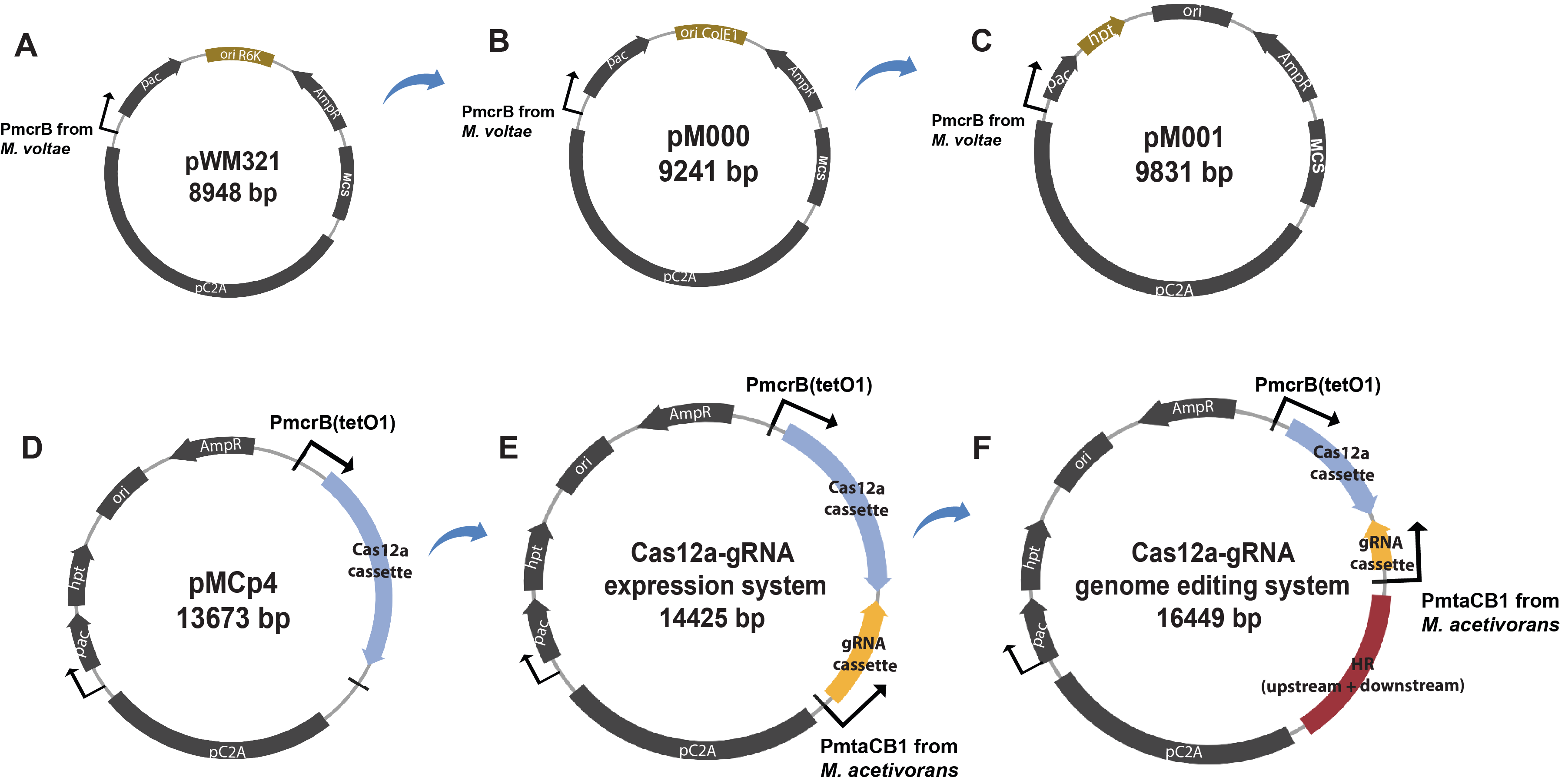

**Supplementary Figure 1. Plasmid maps of the Cas12a-mediated editing system.** **(A)** Plasmid pWM321, *Escherichia coli/Methanosarcina* shuttle vector. Plasmid components are shown. pC2A, a naturally occurring plasmid from *Methanosarcina acetivorans*. *pac*, puromycin acetyltransferase, which is regulated by the promoter from the *Methanococcus voltae* methyl reductase operon (PmcrB). *ori* R6K, the origin of replication from plasmid R6K enables the plasmid cloning in *E.coli* strains. AmpR, β-lactamase (*bla*) gene, which confers resistance to ampicillin. MCS, multiple cloning site. **(B)** Plasmid pM000, pWM321-derived plasmid where the *ori* R6K is replaced by the *ori* from plasmid ColE1 (*ori* ColE1). **(C)** Plasmid pM001, pM001-derived plasmid where the hypoxanthine phosphoribosyltransferase (*hpt*) gene was inserted downstream of *pac* and driven by PmcrB. **(D)** Plasmid pMCp4 expressing the Cas12a protein in *M. acetivorans*, where the MCS region from pM001 was replaced by the Cas12a cassette. Cas12a cassette consists of the Cas12a gene from *Lachnospiraceae bacterium* (Lb) and the promoter PmcrB (tetO1). **(E)** The Cas12a-gRNA expression system generates double-stranded breaks (DSB) in genome. gRNA cassette, the gRNA containing spacer and direct repeat sequences was expressed by the promoter from *M. acetivorans* methanol-specific methyltransferase operon 1 (PmtaCB1). **(F)** The Cas12a-gRNA genome editing system repairs the DSB with homologous repair (HR) arms to facilitate the genome editing in *M. acetivorans.*

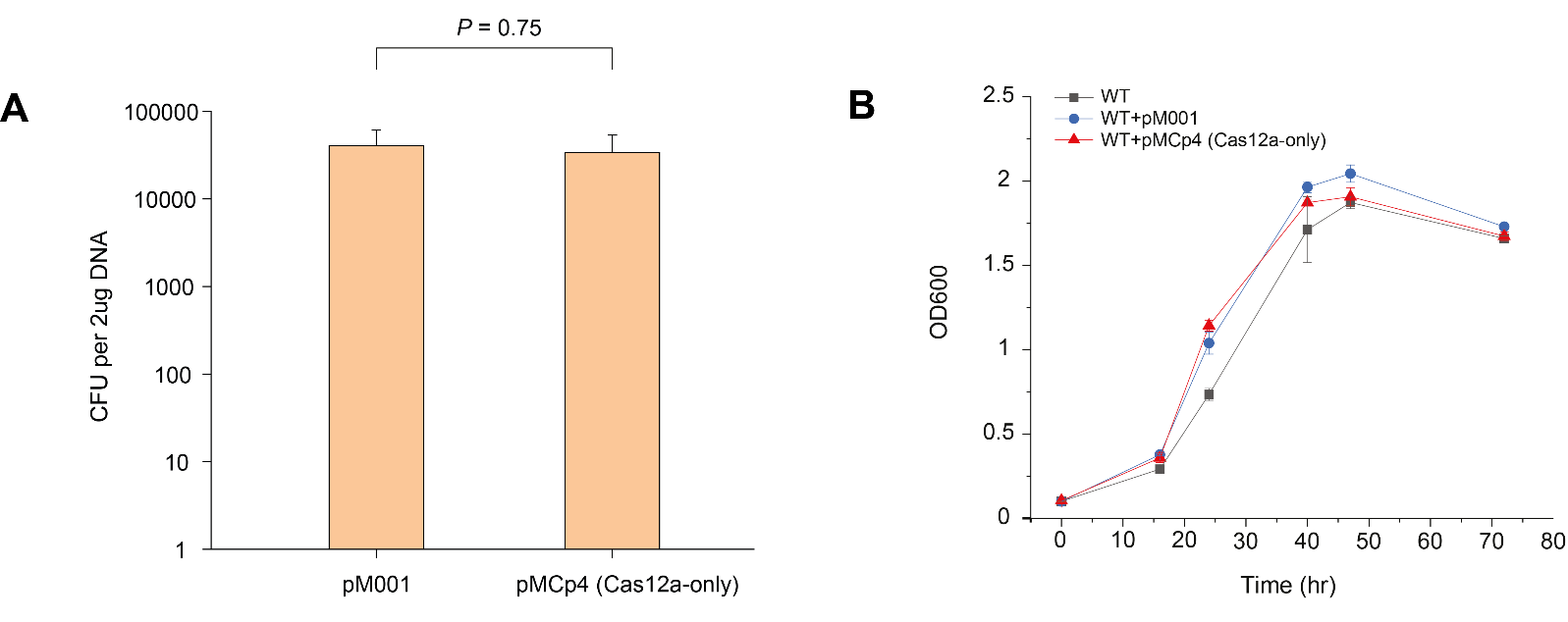

**Supplementary Figure 2. Assessing the toxicity of Cas12a to *M. acetivorans*.** **(A)** Transformation efficiency of the empty vector pM001 and the Cas12a-expressing plasmid pMCp4. Error bar represents the standard deviation of triplicate measurements. *P* = 0.75 (two-tailed *t-test*), indicates no significant difference between the groups. **(B)** Growth curves of *M. acetivorans* cells containing an empty vector or a vector expressing Cas12a. WT, wide type *M. acetivorans* strain. WT+pM001, *M. acetivorans* with empty vector pM001. WT+pMCp4 (Cas12a-only), *M. acetivorans* with Cas12a-expressing plasmid pMCp4. Error bar represents the standard deviation of triplicate experiments.

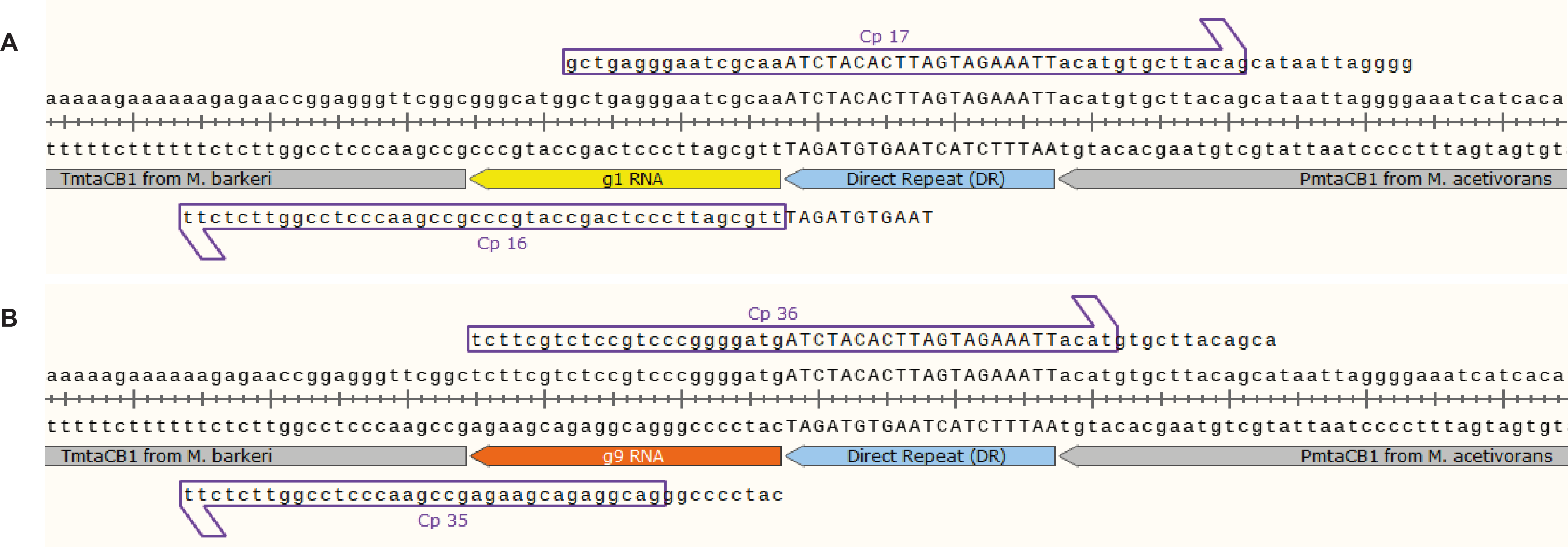

**Supplementary Figure 3. Design of gRNA cassette for Cas12a-gRNA expression system.** **(A)** The g1RNA cassette designed for targeting *ssuC*. The promoter (PmtaCB1) and terminator (TmtaCB1) from the methanol-specific methyltransferase operon in *M. acetivorans* and *M. barkeri* were added separately to the gRNA cassette constructs. g1RNA and direct repeat (DR) sequences were amplified by PCR using primers Cp16 and Cp17. Cp16, forward primer for amplifying terminator TmtaCB1. Cp17, reverse primer for amplifying promoter PmtaCB1. **(B)** The g9RNA cassette designed for targeting *frhA*. g9RNA and direct repeat (DR) sequences were amplified by PCR using primers Cp35 and Cp36. Cp35, forward primer for amplifying terminator TmtaCB1. Cp36, reverse primer for amplifying promoter PmtaCB1.

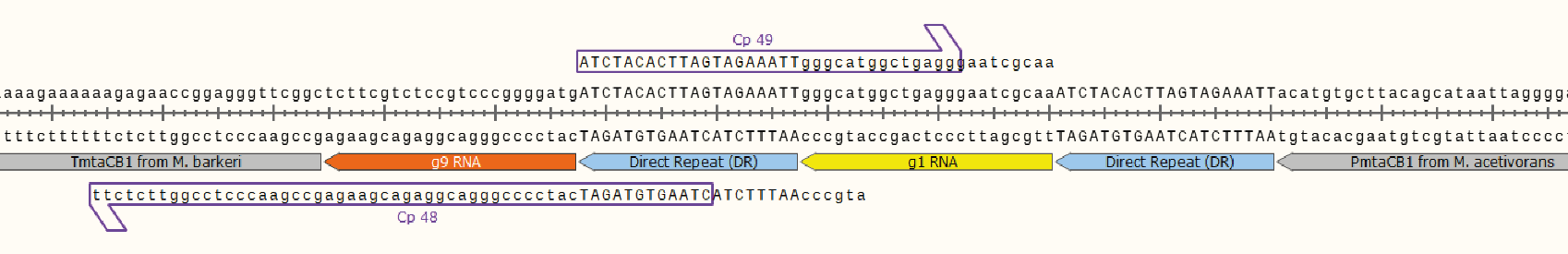

**Supplementary Figure 4. Design of crRNA array for Cas12a-mediated multiplex genome editing.** The crRNA array targeting *ssuC* and *frhA* to simultaneously generate two leakages on the genome. The promoter (PmtaCB1) and terminator (TmtaCB1) from methanol-specific methyltransferase operon in *M. acetivorans* and *M. barkeri* were added to the constructs. g1RNA, g9RNA, and relevant direct repeat (DR) sequences were amplified by PCR using primers Cp48 and Cp49 with the previously constructed pMCp2-g1RNA as the template. Cp48, forward primer for amplifying terminator TmtaCB1. Cp49, reverse primer for amplifying promoter PmtaCB1.

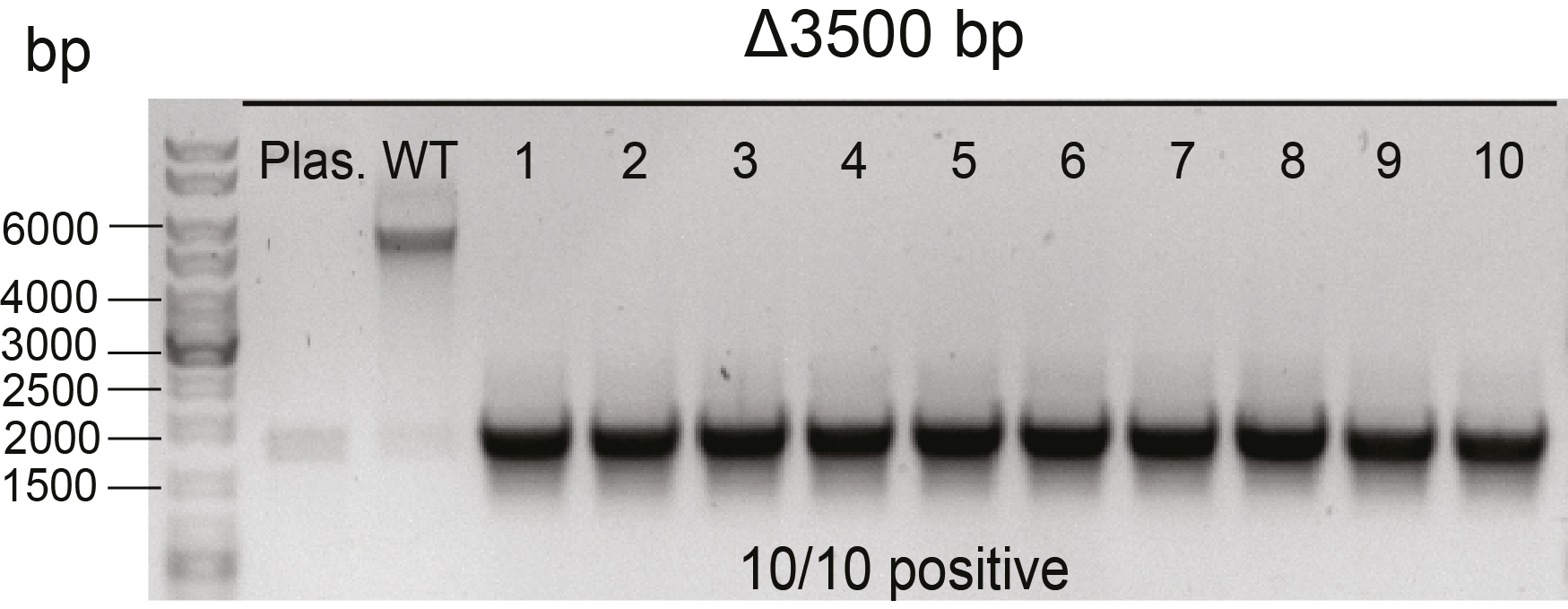

**Supplementary Figure 5. Cas12a-mediated *frhADGB* operon deletion in *M. acetivorans*.** Δ3500 bp, plasmid targeting *frhA* generates a 3500-bp deletion. Plas. and WT, plasmid (pMCp3-g9-3500) and wild-type *M. acetivorans* genome served as the negative control in colony PCR. Thermo Scientific™ GeneRuler DNA ladder mix was used for sizing DNA fragments. 1-10, ten Pur^R^ transformants randomly selected for colony PCR.

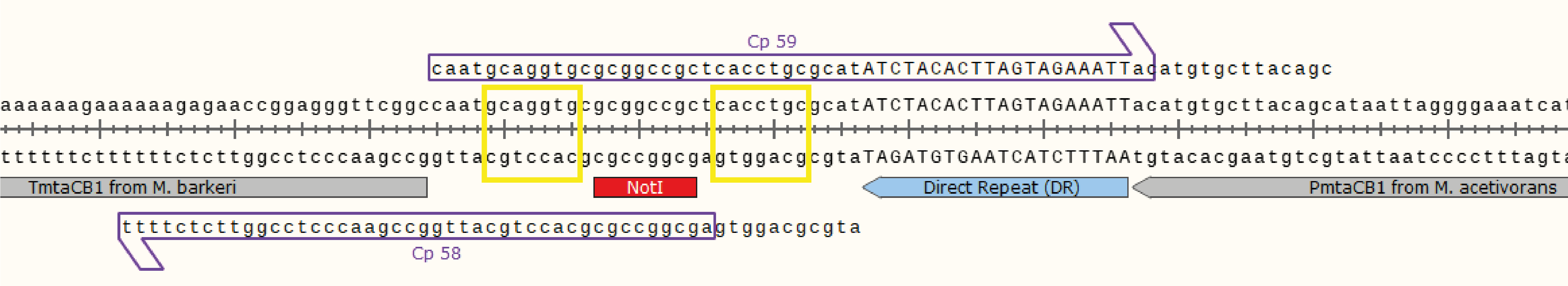

**Supplementary Figure 6. Design of exchangeable gRNA sequences for plasmid pMCp2-gX.** All sites for the Type IIS restriction enzyme *PaqCI (AarI)* were removed from the plasmid pMCp4. The promoter (PmtaCB1) and terminator (TmtaCB1) from methanol-specific methyltransferase operon in *M. acetivorans* and *M. barkeri* were separately added to the gRNA cassette construct. In the gRNA cassette constructs, sites for two *AarI* and one *NotI* were placed between the direct repeat (DR) and the TmtaCB1 sequences by PCR using primers Cp58 and Cp59. *AarI* recognition sequences (5'-CACCTGC-3') are marked by yellow squares. Cp58, the forward primer for amplifying terminator TmtaCB1. Cp59, the reverse primer for amplifying promoter PmtaCB1.

**Easy-to-use outline for cloning with plasmid pMCp2-gX:**

1. Design primers Cp58 and Cp59. The overlapping region between the primers is the gRNA sequences that will be assembled.
2. Anneal primers Cp58 and Cp59 directly with Thermo cycler using an optimal annealing temperature to amplify the overlapping DNA. 5'- and 3'-ends are separately complementary to TmtaCB1 and PmtaCB1.
3. Linearize plasmid pMCp2-gX by digesting with AarI restriction enzyme. NotI restriction enzyme can also be added to increase digestion efficiency (optional).
4. Gibson assembly method is used for assembling together the overlapping DNA product and linearized plasmid pMCp2-gX.
5. Transform the assembled construct into competent *E.coli* cells and cultivate overnight at 37°C. Screen for positive constructs by colony PCR.

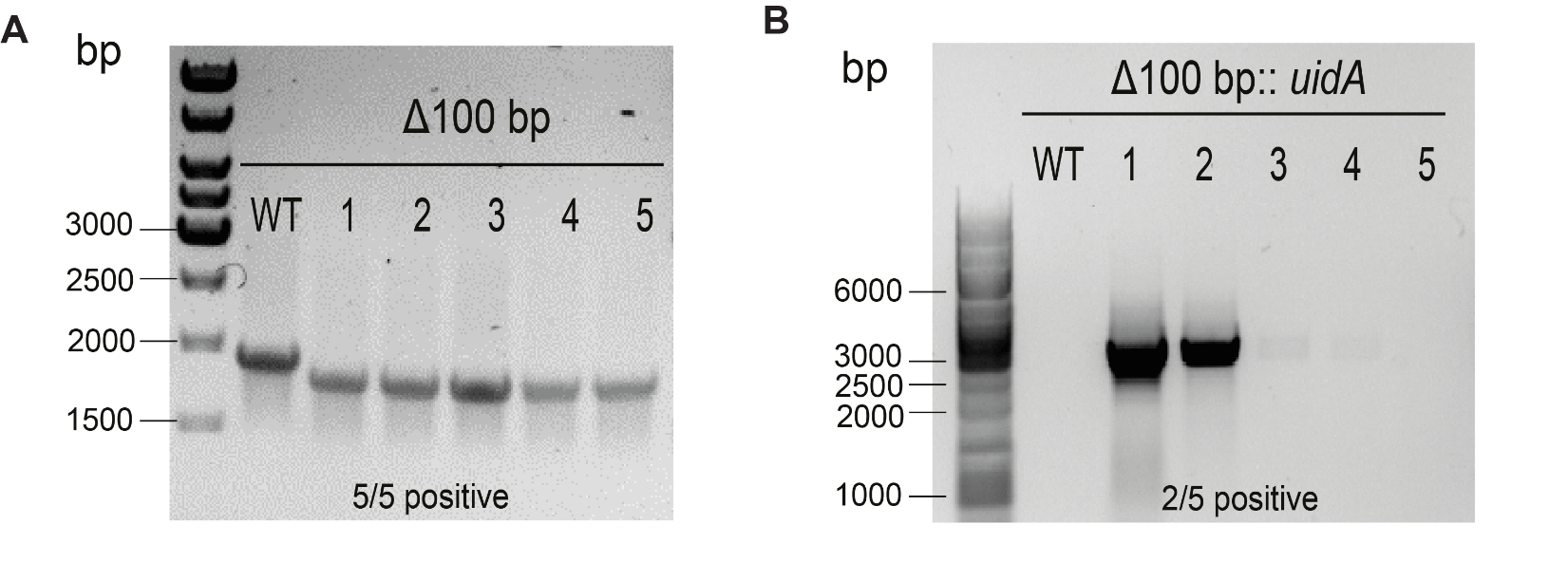

**Supplementary Figure 7. Plasmid curing efficiency after genome editing. (A)** Colony PCR of 8ADP^R^ M73-100 isolates. Δ100 bp, genome-edited *M. acetivorans* transformants with 100-bp deletion in *ssuC*. WT, wild type *M. acetivorans* genome served as the negative control. **(B)** Colony PCR of 8ADP^R^ M73-uid isolates. Δ100 bp::*uidA*, genome-edited *M. acetivorans* transformants with 2400-bp *uidA* cassette insertion in *ssuC*. WT, wild type *M. acetivorans* genome served as the negative control. Thermo Scientific™ GeneRuler 1kb DNA ladder was used for sizing DNA fragments.
