## Supplementary table S3 for "CRISPR/Cas12a toolbox for genomic manipulation in *Methanosarcina acetivorans*"

**Table 3. Primers used in this study**

| **Primer** | **Sequence** | **Source** | **Description** |
| --- | --- | --- | --- |
| Cp1 | ctgaacacgcggcgcagttc | This study | 001-F1-F |
| Cp2 | cttccggctggctggtttattg | This study | 001-F1-R |
| Cp3 | ccggctccagatttatcagc | This study | 001-F2-F |
| Cp4 | taatggagagttgaagtgggaaggac | This study | 001-F2-R |
| Cp5 | catcgttttcctgtccttcccac | This study | 001-F3-F |
| Cp6 | catggggtcgtgcgctcctttc | This study | 001-F3-R |
| Cp7 | gaaaggagcgcacgaccccatg | This study | 001-F4-F |
| Cp8 | gaactgcgccgcgtgttcag | This study | 001-F4-R |
| Cp9 | cggcccacgtggccactagtacttctcgaggcatgcttcatttatcggagaacacaaaag | This study | Pmcr(tetO1)-F |
| Cp10 | atgaatttcctccttaatttattaaaatcattttgggactggtcacctac | This study | Pmcr(tetO1)-R |
| Cp11 | cccaaaatgattttaataaattaaggaggaaattcatatgagcaagctggagaagtttac | This study | LbCas12a-F |
| Cp12 | ggcctactctgttttaaactgttgaatttattgagtttagctggtctgggcgtactc | This study | LbCas12a-R |
| Cp13 | actcaataaattcaacagtttaaaacagagtaggcc | This study | Tmcr-F |
| Cp14 | aacccgggccctatatatggatccaatagaattatatgagcctgtaacggggat | This study | Tmcr-R |
| Cp15 | ccccgttacaggctcatataattctattcctgcagggccttttaaaaagggattgagcgaaaa | This study | TmtaCB1-R |
| Cp16 | taagtgtagatttgcgattccctcagccatgcccgccgaaccctccggttctctt | This study | Tmta-g1-F |
| Cp17 | gctgagggaatcgcaaatctacacttagtagaaattacatgtgcttacagcataattagggg | This study | Pmta-g1-R |
| Cp18 | aattataacccgggccctatatatggatcccctgcaggaacaacatcagtcacctaaaaagagaaaac | This study | PmtaCB1-F |
| Cp19 | gactgatgttgttcctgcaggggatcccgccggcgttagcagttttttctttatcggcttcttca | This study | HR-100up-F |
| Cp20 | aacgtaccgagggtttatgttgagcggccgcccttgtagctgcagagatgttcgg | This study | HR-100up-R |
| Cp21 | tacaagggcggccgctcaacataaaccctcggtacgtt | This study | HR-100dw-F |
| Cp22 | aggtaattataacccgggccctatatatcgccggcggtgaaggcaatggacgttcga | This study | HR-100dw-R |
| Cp23 | ggtgactgatgttgttcctgcaggggatcccgccggcgggatatccgtgtaagacccggatg | This study | HR-500up-F |
| Cp24 | cccctgggaatagctatgggatgggcggccgcatgggtagagtaagtgtaaaaaatgtttctc | This study | HR-500up-R |
| Cp25 | ctctacccatgcggccgcccatcccatagctattcccagggg | This study | HR-500dw-F |
| Cp26 | tgaggtaattataacccgggccctatatatcgccggcggaaagggctcaaaattgccactt | This study | HR-500dw-R |
| Cp27 | gtgactgatgttgttcctgcaggggatcccgccggcgaaaaaggagcttttctcgaagaagacc | This study | HR-1000up-F |
| Cp28 | ctaagaatccttttaattagacatgaaaattaatttcacaaagcggccgctccagcagtattctctctttccctgga | This study | HR-1000up-R |
| Cp29 | actgctggagcggccgctttgtgaaattaattttcatgtctaattaaaaggattcttag | This study | HR-1000dw-F |
| Cp30 | tgaggtaattataacccgggccctatatatcgccggcgagaaaggatggtggcaggaag | This study | HR-1000dw-R |
| Cp31 | gtgactgatgttgttcctgcaggggatcccgccggcgcatggaagataatctggccttctttgc | This study | HR-2000up-F |
| Cp32 | tcttccggagaaatggaagcaaacgcggccgccggttcagcaagtgaaatgct | This study | HR-2000up-R |
| Cp33 | tgaaccggcggccgcgtttgcttccatttctccggaaga | This study | HR-2000dw-F |
| Cp34 | tgaggtaattataacccgggccctatatatcgccggcgagatgtattaatacatttgggtcataatagaaaagcat | This study | HR-2000dw-R |
| Cp35 | catccccgggacggagacgaagagccgaaccctccggttctctt | This study | Tmta-g9-F |
| Cp36 | tcttcgtctccgtcccggggatgatctacacttagtagaaattacatgtgcttacagca | This study | Pmta-g9-R |
| Cp37 | ggtgactgatgttgttcctgcaggggatcctcaacaaacggtgtttcagcactgg | This study | HR-3500up-F |
| Cp38 | gaaaaaaaggcggccgccatcccctcgctttcgattctgtac | This study | HR-3500up-R |
| Cp39 | ggatggcggccgcctttttttcgtcatagaaattattcggcgttattgc | This study | HR-3500dw-F |
| Cp40 | aattataacccgggccctatatatatagttgcttggaatcttgtagttggtatgt | This study | HR-3500dw-R |
| Cp41 | gaacatctctgcagctacaaggaatatcatttcgtcattttcctaagaaa | This study | uidA cas-Pmcr-F |
| Cp42 | ggggtttctacaggacgtaacattttaatttcctccttaatttattaaaatcattttgggact | This study | uidA cas-Pmcr-R |
| Cp43 | agtcccaaaatgattttaataaattaaggaggaaattaaaatgttacgtcctgtagaaacccca | This study | uidA cas-uidA-F |
| Cp44 | ggcctactctgttttagtatgttcatttattgagttcattgtttgcctccctgctg | This study | uidA cas-uidA-R |
| Cp45 | cagcagggaggcaaacaatgaactcaataaatgaacatactaaaacagagtaggcc | This study | uidA cas-Tmcr-F |
| Cp46 | ggttttaaaaacgtaccgagggtttatgttgagggagaattatatgagcttataacggtagaaatattgttt | This study | uidA cas-Tmcr-R |
| Cp47 | tcaacataaaccctcggtacgtt | This study | uidA cas-HR-100dw-F |
| Cp48 | atgcccaatttctactaagtgtagatcatccccgggacggagacgaagagccgaaccctccggttctctt | This study | Tmta-g1g9-F |
| Cp49 | atctacacttagtagaaattgggcatggctgagggaatcgcaa | This study | Pmta-g1g9-R |
| Cp51 | tgttgttcctgcaggggatccgcactcactttggcttctgggttgcc | This study | HR-g9-100up-F |
| Cp52 | ctataagcggccgcaggatcaggatattgtgcagggcg | This study | HR-g9-100up-R |
| Cp53 | ctgatcctgcggccgcttatagccgtcgcaggcggcgag | This study | HR-g9-100dw-F |
| Cp54 | aattataacccgggccctatatatgcgggtccgaacccatcgtccgc | This study | HR-g9-100dw-R |
| Cp55 | gtgaaggcaatggacgttcgattgtatc | This study | g1HR-100dw-R |
| Cp56 | atcgaacgtccattgccttcacgcactcactttggcttctgggttgc | This study | g9HR-100up-F |
| Cp57 | gtaattataacccgggccctatatatcgccggcggcgggtccgaacccatcgtccgc | This study | g9HR-100dw-R |
| Cp58 | atgcgcaggtgagcggccgcgcacctgcattggccgaaccctccggttctctttt | This study | Tmta-gX-F |
| Cp59 | caatgcaggtgcgcggccgctcacctgcgcatatctacacttagtagaaattacatgtgcttacagc | This study | Pmta-gX-R |
| **Colony PCR primer** | **Sequence** | **Source** | **Description** |
| veri1 | ccacaagctccaggtatttttcca | This study | Δ100-F |
| veri2 | gaaagggctcaaaattgccactttccc | This study | Δ100-R |
| veri3 | aaaaaggagcttttctcgaagaagacctt | This study | Δ500-F |
| veri4 | agaaaggatggtggcaggaagatcttg | This study | Δ500-R |
| veri5 | ggatatccgtgtaagacccggatgcac | This study | Δ1000/Δ2000-F |
| veri6 | cagattgcatacatgactgcctatgag | This study | Δ1000-R |
| veri7 | cccttattgtgacaagcctgccagcg | This study | Δ2000-R |
| veri8 | tcccgccgggaatggtgattacc | This study | veri uidA-F |
| veri9 | aaagaaggctatcctcgaattctct | This study | Δ3500-F |
| veri10 | ctcaaatcgttagaccagaaacac | This study | Δ3500-R |
| veri11 | tgccgaagagattttcagcaagaatgtgag | This study | genome frh dele-F |

Note: gRNA sequences are underlined.
